# Comprehensive, integrated, and phased whole-genome analysis of the primary ENCODE cell line K562

**DOI:** 10.1101/192344

**Authors:** Bo Zhou, Steve S. Ho, Stephanie U. Greer, Xiaowei Zhu, John M. Bell, Joseph G. Arthur, Noah Spies, Xianglong Zhang, Seunggyu Byeon, Reenal Pattni, Noa Ben-Efraim, Michael S. Haney, Rajini R. Haraksingh, Hanlee P. Ji, Giltae Song, Dimitri Perrin, Wing H. Wong, Alexej Abyzov, Alexander E. Urban

## Abstract

K562 is widely used in biomedical research. It is one of three tier-one cell lines of ENCODE and also most commonly used for large-scale CRISPR/Cas9 screens. Although its functional genomic and epigenomic characteristics have been extensively studied, its genome sequence and genomic structural features have never been comprehensively analyzed. Such information is essential for the correct interpretation and understanding of the vast troves of existing functional genomics and epigenomics data for K562. We performed and integrated deep-coverage whole-genome (short-insert), mate-pair, and linked-read sequencing as well as karyotyping and array CGH analysis to identify a wide spectrum of genome characteristics in K562: copy numbers (CN) of aneuploid chromosome segments at high-resolution, SNVs and Indels (both corrected for CN in aneuploid regions), loss of heterozygosity, mega-base-scale phased haplotypes often spanning entire chromosome arms, structural variants (SVs) including small and large-scale complex SVs and non-reference retrotransposon insertions. Many SVs were phased, assembled, and experimentally validated. We identified multiple allele-specific deletions and duplications within the tumor suppressor gene *FHIT*. Taking aneuploidy into account, we re-analyzed K562 RNA-seq and whole-genome bisulfite sequencing data for allele-specific expression and allele-specific DNA methylation. We also show examples of how deeper insights into regulatory complexity are gained by integrating genomic variant information and structural context with functional genomics and epigenomics data. Furthermore, using K562 haplotype information, we produced an allele-specific CRISPR targeting map. This comprehensive whole-genome analysis serves as a resource for future studies that utilize K562 as well as a framework for the analysis of other cancer genomes.

## INTRODUCTION

K562 is an immortalized chronic myelogenous leukemia (CML) cell line derived from a 53-year-old Caucasian female in 1970 (Lozzio and Lozzio 1975). Since being established, K562 has been widely used in biomedical research as a “work-horse” cell line, resulting in over 17,000 publications to date. In most cases, its use is similar to that of a model organism, contributing to the understanding of fundamental human biological processes as well as to basic and translational cancer research (Grzanka et al. 2003; Drexler et al. 2004; Butler and Hirano 2014). Along with the H1 human embryonic stem cell line and the GM12878 lymphoblastoid cell line, K562 is one of the three tier-one cell lines of the ENCyclopedia Of DNA Elements Project (ENCODE) (The ENCODE Project Consortium 2012), forming the basis of over 1,300 ENCODE datasets to date (Sloan et al. 2016). Furthermore, it is also one of the few cell lines most commonly used for large-scale CRISPR/Cas9 gene-targeting screens (Wang et al. 2015; Arroyo et al. 2016; Morgens et al. 2016; Han et al. 2017; Adamson et al. 2016; Liu et al. 2017).

Although the functional genomic characteristics of K562 have been extensively studied and documented, reflected in close to 600 ChIP-seq, 400 RNA-seq, 50 DNase-Seq, and 30 RIP-Seq datasets available through the ENCODE portal (Sloan et al. 2016), the sequence and structural features of the K562 genome have never been comprehensively characterized, even though past cytogenetic studies using G-banding, fluorescence *in situ* hybridization (FISH), multiplex-FISH, and comparative genomic hybridization (CGH) showed that K562 cells contain pervasive aneuploidy and multiple gross structural abnormalities (Selden et al. 1983; Wu et al. 1995; Naumann et al. 2001; Gribble et al. 2000), not unexpected for a cancer cell line. In other words, the rich amount of K562 functional genomics and epigenomics work conducted to date, in particular integrative analyses that have been carried out in various settings using the vast troves of K562 ENCODE data, were done without taking into account the many differences of the K562 genome relative to the human reference genome. This leads to skewed interpretations and reduces the amount of knowledge that can be gained from the rich, multi-layered ENCODE datasets that continue to accumulate.

Here, we report for the first time a comprehensive characterization of the K562 genome that include copy numbers (CN) of chromosome segments at high-resolution, single-nucleotide variants (SNVs, also including single-nucleotide polymorphisms, i.e. SNPs) and small insertions and deletions (Indels) with allele-frequencies corrected by CN in aneuploid regions, loss of heterozygosity, mega-base-scale phased haplotypes often spanning entire chromosome arms, and structural variants (SVs) including small and large-scale complex SVs with phasing. We then took first steps into exploring how knowledge about genome sequence and structural features can influence the interpretation of functional genomics and epigenomics data and show examples of how deeper insights into genome regulatory complexity can be obtained by integrating genomic context. These insights also shed light on important questions regarding cancer evolution.

## RESULTS

### Karyotyping

The K562 cell line exhibits pervasive aneuploidy (Fig. 2A). Analysis of 20 individual K562 cells using GTW banding showed that all cells demonstrated a near-triploid karyotype and are characterized by multiple structural abnormalities. The karyotype of our line of K562 cells is overall consistent (although not identical) with previously published karyotypes (Selden et al. 1983; Wu et al. 1995; Naumann et al. 2001; Gribble et al. 2000), suggesting that its near-triploid state arose during leukemogenesis or early in the establishment of the cell line. It also suggests that different K562 cell lines kept and passaged in different laboratories may exhibit some additional karyotypic differences. Although the karyotype for all chromosomes in our K562 cell line was supported by previous karyotype analyses, slight variations do exist among the various published analyses (Supplemental Table S1) with chromosomes 10, 12, and 21 showing the most variability.

### Identification of Copy Number (CN) by Chromosome Segments

We used read-depth analysis (Abyzov et al. 2011) to assign a CN i.e. ploidy to all chromosome segments at 10kb-resolution or entire chromosomes in the K562 genome (Fig. 1, Supplemental Table S2). We first calculated WGS coverage in 10 kb bins across the genome and plotted it against %GC content where five distinct clusters were clearly observed (Supplemental Fig. S2). Clusters were designated as corresponding to particular CNs based on the mean coverage of each cluster (Supplemental Methods). Such designations confirm that the triploid state is the most common in the K562 genome. The CN assigned to all chromosome segments using this approach are consistent with array CGH (Supplemental Fig. S3, Supplemental Data) and also with previous CGH analyses (Gribble et al. 2000; Naumann et al. 2001) with minor differences on chromosomes 7, 10, 11, and 20 (Supplemental Table S3). While on a general level, the CNs identified based on read-depth analysis tracks the findings from karyotyping, read-depth analysis reveals the CNs of many chromosome segments that would not have been apparent from karyotyping alone (Supplemental Fig. S3, Supplemental Data, Supplemental Table S2). We see that 53.5% of the K562 genome has a baseline CN of three (consistent with the karyotype, Fig. 2A), 16.9% CN of four, 1.9% CN of five, 2.4% CN of one, and only 30.0% has remained in a diploid state (Figure 2B). In addition, two large regions (5.8 Mb and 3.1 Mb in size) on chromosome 9 (20,750,000-26,590,000 and 28,560,00031,620,000 respectively) were lost entirely (Supplemental Table S2).

**Figure 1.**
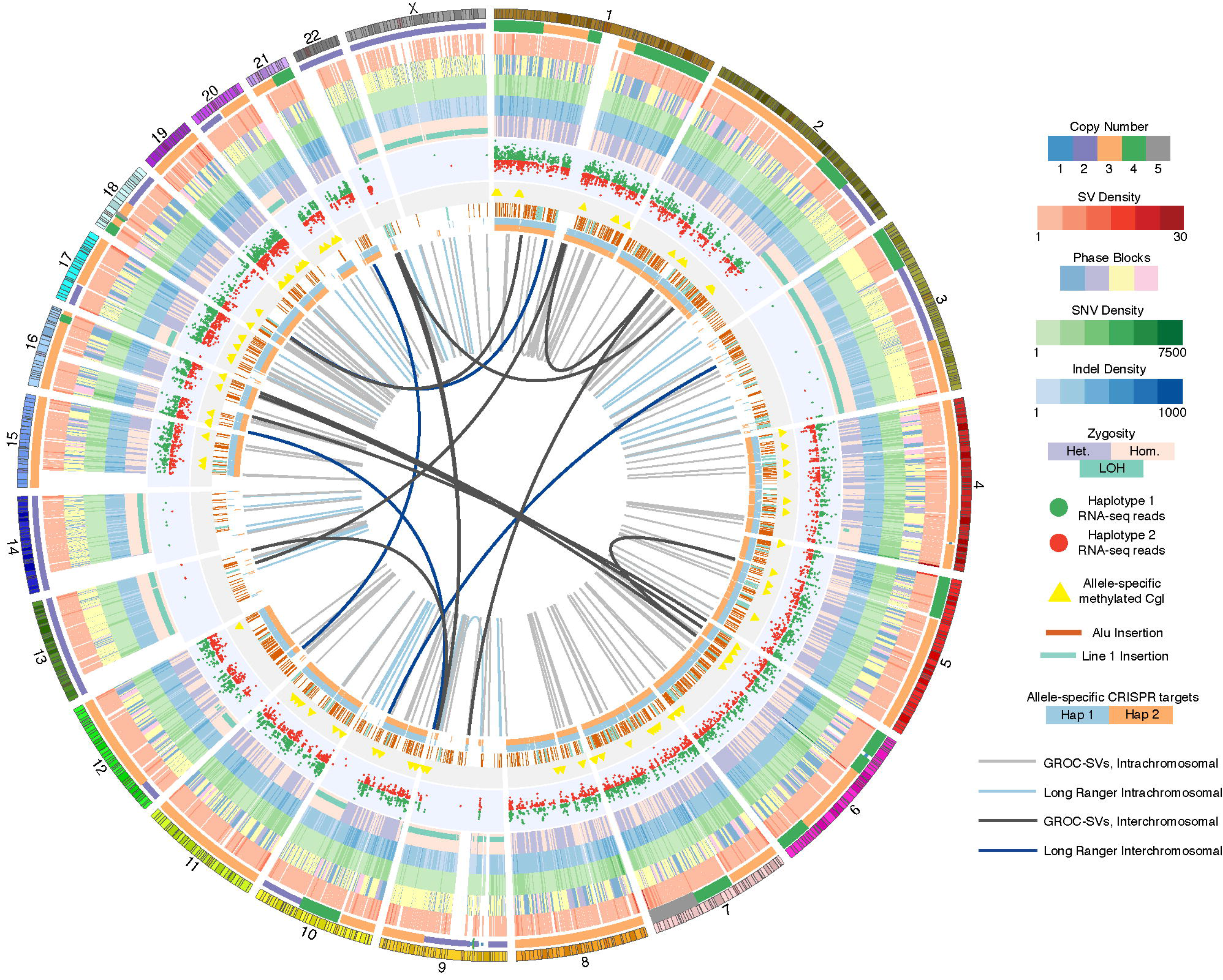
Comprehensive overview of the K562 genome. Circos (Krzywinski et al. 2009) visualization of K562 genome with the following tracks in inward concentric order: chromosomes; CN i.e. ploidy by chromosome segment; merged SV density in 1.5 Mb windows of deletions, duplications, and inversions identified using ARC-SV (Arthur et al. 2017), BreakDancer (Chen et al. 2009), BreakSeq (Lam et al. 2010), LUMPY (Layer et al. 2014), Pindel (Ye et al. 2009), and Long Ranger (Zheng et al. 2016; Marks et al. 2018); phased haplotype blocks (demarcated with 4 colors for clearer visualization); SNV density in 1 Mb windows; Indel density in 1 Mb windows; dominant zygosity in 1 Mb windows (heterozygous or homozygous > 50%) with regions exhibiting loss of heterozygosity (LOH) indicated; RNA-seq reads for loci exhibiting allele-specific expression; CpG islands (CgI) exhibiting allele-specific methylation; histogram (log-scale) of allele-specifically methylated CpGs in 50 kb windows; nonreference Alu and LINE-1 insertions; allele-specific CRISPR target sites; large-scale rearrangements detected by Long Ranger (Zheng et al. 2016; Marks et al. 2018) (light blue: intrachromosomal: dark blue: interchromosomal); and by GROC-SVs (Spies et al. 2017) (light-gray: intrachromsomal; dark-gray: interchromosomal).

**Figure 2.**
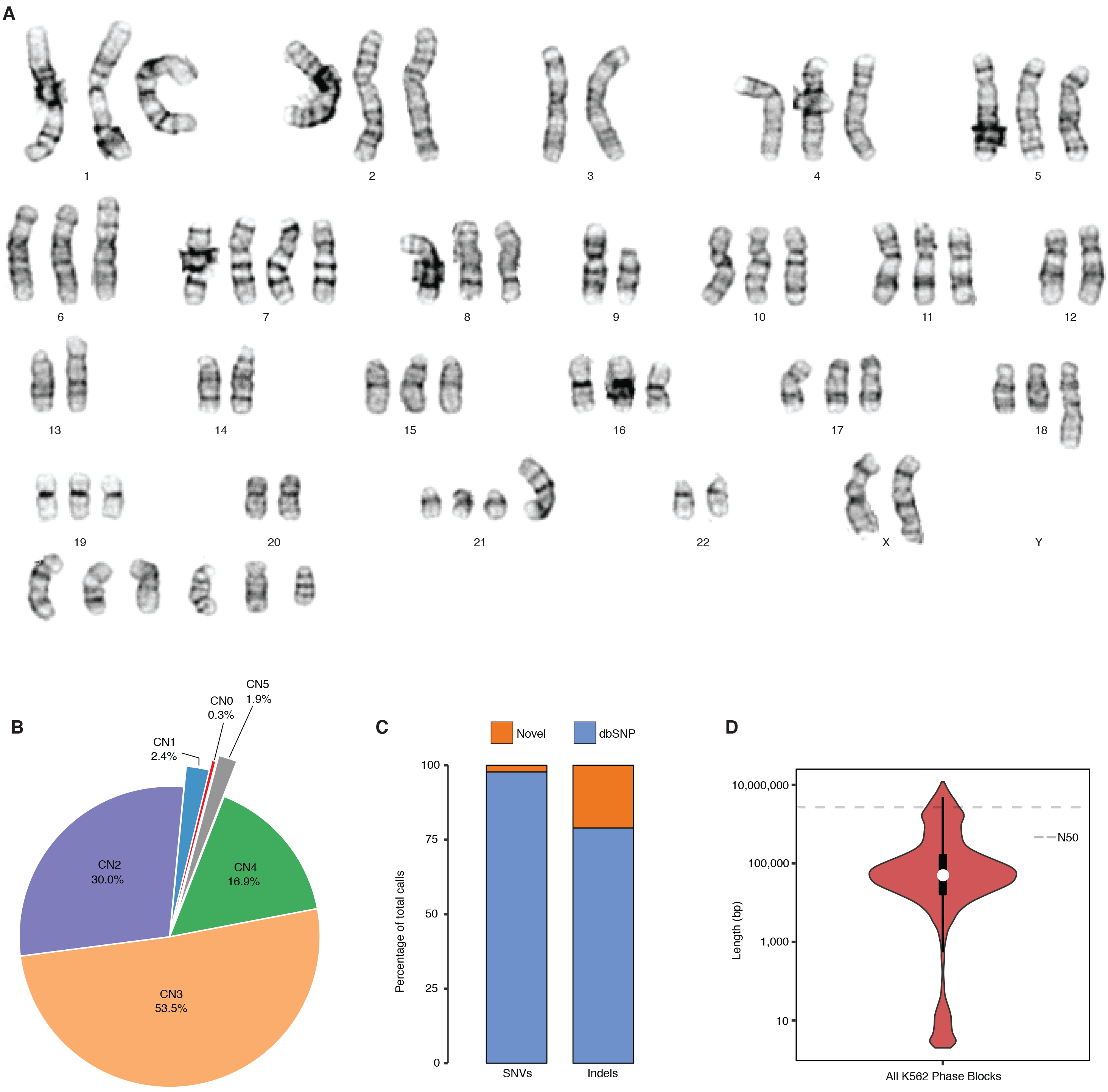
K562 ploidy and haplotypes. (*A*) Representative karyogram of K562 cells produced by GTW banding showing multiple numerical and structural chromosomal abnormalities and an overall near triploid karyotype. ISCN 2013 description in relationship to a triploid karyotype [<3n>]: 53~70<3n>,XX,-X or Y, −3,?dup(6)(p21p25),+7,?inv(7)(p13p22),add(7)(q32),−9,add(9)(p24),del(9)(p13),add(10)(q22),−13,add(13)(p11),−14,add(17)(p11.2)×2,add(18)(q23),−20,der(21)t(1;21)(q21;p11),−22,+4~7mar[cp20]. (*B*) CN (i.e. ploidy) by percentage across the K562 genome. (*C*) Percentage of K562 SNVs and Indels that are novel and known in dbSNP (Sherry et al. 2001). (*D*) Violin plot, with overlaid boxplot, of phased haplotype block sizes (Y-axis, log-scaled) where the dashed line represents the N50 value (2,721,866 bp).

### SNVs and Indels

We identified SNVs and Indels in the K562 genome. By taking into account the CN of the chromosomal segments in which they reside, we assigned heterozygous allele frequencies to these variants, including non-conventional frequencies (e.g. 0.33 and 0.67 in triploid regions; 0.25, 0.50, and 0.75 in tetraploid regions). Using this approach, we detected and genotyped a total of 3.09 M SNVs (1.45 M heterozygous, 1.64 M homozygous) and 0.70 M Indels (0.39 M heterozygous, 0.31 M homozygous) (Table 1, Dataset S1). Interestingly, there are 13,471 heterozygous SNVs and Indels that have more than two haplotypes in aneuploid regions where CN is >2 (Dataset S1). Furthermore, chromosomes 3, 9, 13, 14, and X along with large stretches of chromosomes 2, 10, 12, 17, 20, and 22 show striking loss of heterozygosity (LOH) (Fig. 1 and Supplemental Table S4). While a normal tissue sample corresponding to K562 is not available for comparative analysis, we overlapped these SNVs and Indels with those in dbsnp138 (Sherry et al. 2001) and found the overlap to be 98% and 79% respectively (Fig. 2C, Dataset S1), suggesting an accumulation of a significant number of K562-specific SNVs and Indels relative to germline variants present in the population. After filtering for protein-altering SNVs and Indels in K562 that overlap with those identified from the 1000 Genomes Project or from the Exome Sequencing Project, we found that 424 SNVs and 148 Indels are private protein-altering (PPA) (Table 1, Supplemental Table S5). Furthermore, the overlap between the PPA variants and the Catalogue of Somatic Mutations in Cancer (COSMIC) is 53% and 31% for SNVs and Indels respectively (Supplemental Table S6). Eighteen genes that acquired PPA variants overlap with the Sanger Cancer Gene Census; canonical tumor suppressor genes and oncogenes such as *RAD51B*, *TP53*, *PDGFRA*, *RABEP1*, *EPAS1*, and *WHISC1* are notably present among them (Supplemental Table S7).

### Haplotype Phasing

We performed haplotype phasing for the K562 genome by performing 10× Genomics linked-read library preparation and sequencing (Zheng et al. 2016; Marks et al. 2018). This library was sequenced (2 × 151 bp) to 59× genome coverage. Post sequencing quality-control analysis showed that 1.06 ng, or approximately 320 genome equivalents, of high molecular weight (HMW) K562 genomic DNA fragments (average fragment size = 59 kb, 95.3% >20kb, 11.9% >100 kb) were partitioned into 1.56 million oil droplets for uniquely barcoding (16 bp) within each droplet. Half of all reads come from HMW DNA molecules with at least 64 linked-reads (N50 Linked-Reads per Molecule or LPM) (Table 1). We estimate the actual physical coverage (C_F_) to be 191×. The overall sequencing coverage is C = C_R_ × C_F_ = 59×. The length of sequence coverage per 2 × 151 bp paired-ended read minus 16 bp of “HWM fragment barcode” is 286 bp, thus coverage (C_R_) of the average input HMW genomic DNA (59 kb) is 18,304 bp (286 bp × 64 linked-reads) or 31.0% of 50 kb. Using Long Ranger (Marks et al. 2018), 1.41 M (97.2%) of heterozygous SNVs and 0.58 M (83.7%) of Indels (previously identified, Dataset S1) were successfully phased into 4,987 haplotype blocks (Fig. 1, Table 1, and Dataset S2). The longest is 11.95 Mb (N50 = 2.72 Mb) (Fig. 2D, Table 1, and Dataset S2); however, haplotype block lengths vary widely across different chromosomes (Supplemental Fig. S4, Fig. 1) with poorly phased regions corresponding to regions with LOH (Fig. 1, Supplemental Table S4, Dataset S2).

### Mega-Haplotypes Encompassing Entire Chromosome Arms

Leveraging the haplotype imbalance in aneuploid regions, we constructed mega-haplotypes (Table 2, Supplemental Data), often encompassing entire K562 chromosome arms, by “stitching” the phased haplotype blocks derived from linked-reads using a recently published method (Bell et al. 2017). Briefly, we counted the number of linked-read barcodes for each phased heterozygous SNVs assigned to haplotype blocks that contain ≥100 phased SNVs (Dataset S2). Since each barcode is specific to a given HMW DNA molecule, the total number of unique barcodes is directly associated with the number of individual HMW DNA molecules sequenced. In other words, the counting of unique barcode associated with a particular sequence gives the fractional representation of that sequence (or genomic locus). Thus, for each phased haplotype in aneuploid regions with CN>2, major and minor haplotypes can be assigned according to the number of barcodes associated with each haplotype (Fig. 3), where the major haplotype simply has more associated unique barcodes than the minor. In diploid regions, the two haplotypes are expected to have similar barcode counts. A matched normal control genome is required in order to confidently discriminate between the major and minor haplotypes in a case genome (Bell et al. 2017). Because K562 has no matching normal sample, we used a female genome (NA12878) for which linked-read data is publicly available and which is of the same ethnicity as K562 (Fig. 3). After verifying aneuploidy (or haplotype imbalance) by barcode counting and performing the normalization procedures and statistical tests as described in (Bell et al. 2017), we then “stitched” together contiguous phased haplotype blocks based on the imbalance between the major and minor haplotypes. Using this approach, a total of 31 autosomal mega-haplotypes were constructed (Table 2, Supplemental Data); 15 of which encompass entire (or >95%) chromosome arms such as 19p, 19q, 10p, 7p, and 5q (Fig. 3). The average mega-haplotype is 50.7 Mb or approximately 4 times longer than the longest phased haplotype block from Long Ranger (Fig. 2D, Table 1, Dataset S2, Table 2). The longest mega-haplotype is approximately 137 Mb long (4q).

**Figure 3.**
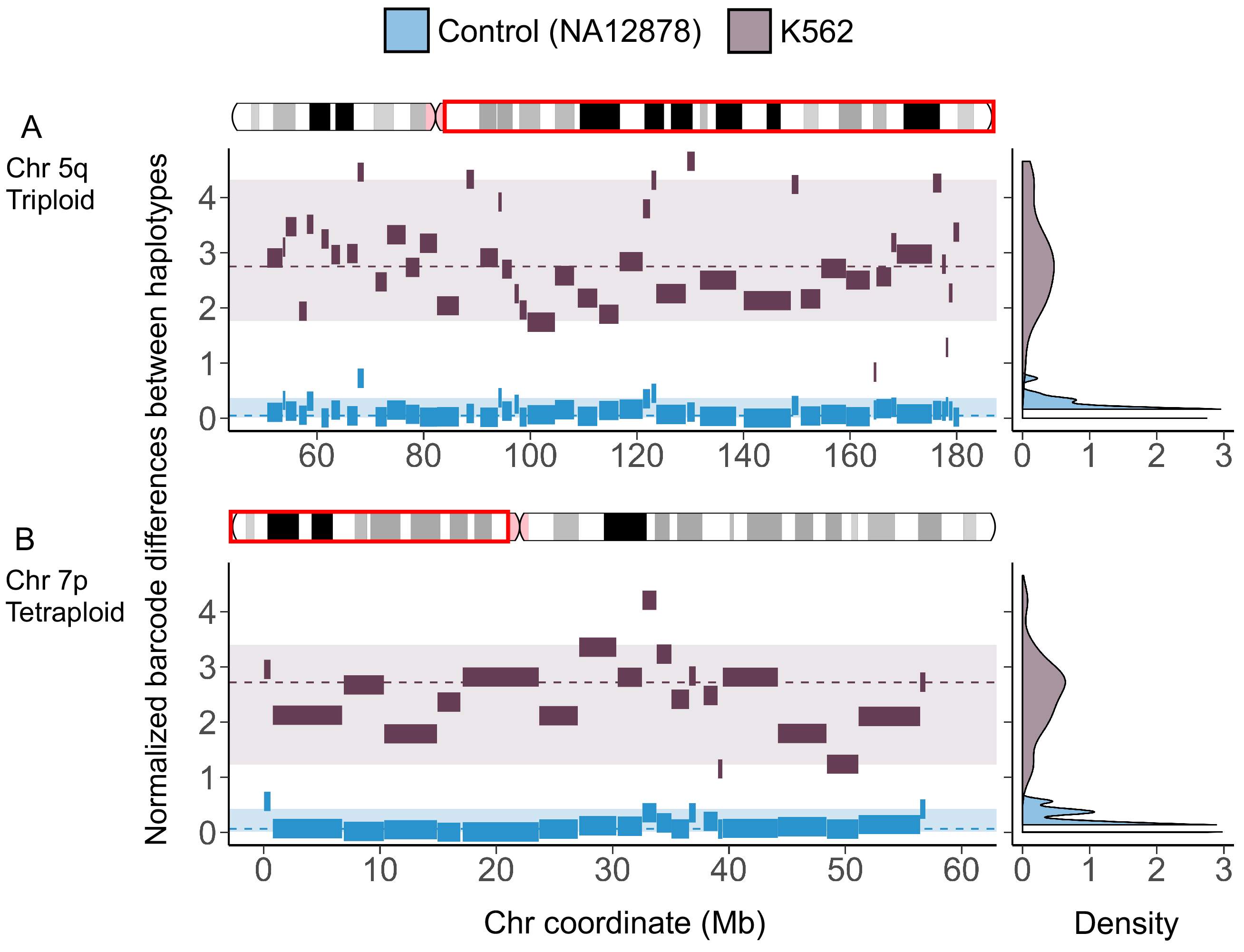
Mega-Haplotypes of entire K562 chromosome arms. X-axis: chromosome coordinate (Mb). Y-axis: difference in unique linked-read barcode counts between major and minor haplotypes, normalized for SNV density. Haplotype blocks from of normal control sample (NA12878) in blue and from K562 in dark gray. Density plots on the right reflects the distribution of the differences in haplotype-specific barcode counts for control sample (blue) and K562 (dark gray). These density distributions are used for testing of significant difference (p<0.001) using one-sided *t*-test. Significant difference in haplotype-specific barcode counts indicate aneuploidy and haplotype imbalance. Haplotype blocks (with ≥ 100 phased SNVs) generated from Long Ranger (Dataset S2) for the major and minor haplotypes were then “stitched” to mega-haplotypes encompassing the entire chromosome arm of (*A*) 5q (triploid) and (*B*) 7p (tetraploid).

### Identification and Reconstruction of Structural Variants (SVs) from Linked-Reads

In addition to phasing, another use for the linked-read sequencing data is to identify breakpoints of large-scale SVs by searching for the discordant mapping of clusters of linked-reads carrying the same barcodes. The identified SVs can then also be assigned to specific haplotypes if the breakpoint-supporting reads contain phased SNVs or Indels (Zheng et al. 2016). Using this approach, which is also implemented by the Long Ranger software from 10× Genomics, we identified 186 large SVs >30 kb (98% phased) (Dataset S3) and 3,541 deletions between 50 bp and 30 kb (79% phased) (Dataset S4). The large SVs include deletions, inversions, duplications, and inter- and intra-chromosomal rearrangements (Dataset S3, Fig. 4A). As expected, we detected the *BCR/ABL1* gene fusion, a hallmark of K562, as one of the SV calls with highest quality score (Fig. 4A, Dataset S3), along with two other known gene fusions in K562 (Engreitz et al. 2012): *XKR3/NUP214* between chromosomes 9 and 22 (Fig. 4A) and *CDC25A/GRID1* between chromosomes 3 and 10 (Dataset S5, Supplemental Data).

**Figure 4.**
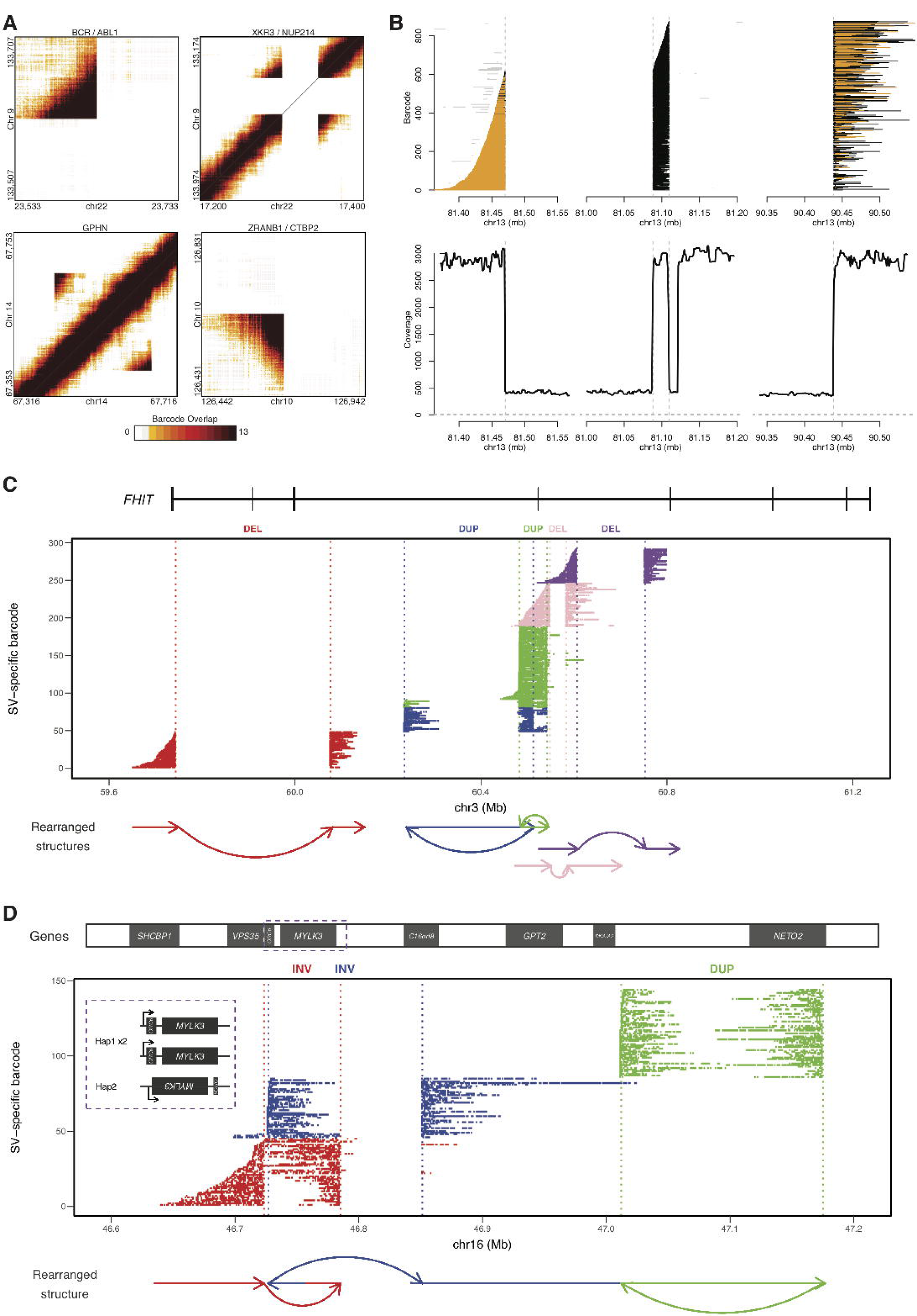
K562 SVs including large complex rearrangements resolved using linked-read sequencing. (*A*) Heat maps of overlapping barcodes for SVs in K562 resolved from linked-read sequencing using Long Ranger (Zheng et al. 2016; Marks et al. 2018). *BCR/ABL1* translocation between chromosomes 9 and 22. *XKR3/NUP214* translocation between chromosomes 9 and 22. Duplication within *GPHN* on chromosome 14. Deletion that partially overlaps *ZRANB1* and *CTB2* on chromosome 10. (*B*) Large complex rearrangement occurring on chromosome 13 with informative reads from only one haplotype (region with loss-of-heterozygosity). Each line depicts a fragment inferred from linked-reads based on clustering of identical barcodes (Y-axis) using GROC-SVs (Spies et al. 2017). Abrupt endings (vertical dased lines) of fragments indicate locations of breakpoints of this complex rearrangement. Fragments are phased locally with respect to surrounding SNVs (colored orange for same haplotype and black when no informative SNVs are found nearby). Gray lines indicate portions of fragments that do not support the current breakpoint. Fragments end abruptly at 81.47 Mb, indicating a breakpoint, picking up again at 81.09 Mb and continuing to 81.11 Mb where they end abruptly, then picking up again at 90.44 Mb. Coverage from 81.12 Mb to 81.20 Mb are from reads with different sets of linked-read barcodes and thus not part of this fragment set. (*C*, *D*) Complex rearrangements involving multiple haplotype-resolved SVs. Using gemtools (Greer et al. 2017), each SV is identified from linked-reads grouped by identical barcodes (i.e. SV-specific barcodes, Y-axis) indicative of single HMW DNA molecules (depicted by each row) that span the breakpoints. SVs are represented in different colors. X-axis: hg19 genomic coordinate. Dotted lines represent individual breakpoints with schematic diagram of the rearranged structures drawn below the plot. (*C*) Multiple SVs within *FHIT* on 3p14.2. (red) Deletion (DEL) (59.74 Mb - 60.08 Mb) results in the loss of multiple exons. Two overlapping duplications (DUP) (blue & green) - the presence of HMW molecules spanning both DUPs indicates a *cis* orientation (same allele of *FHIT)*. Two adjacent DELs (pink & purple) - the spanning HMW molecules for each DEL do not share SV-specific barcodes, indicating that these DELs are in *trans* (different alleles of *FHIT*). SV haplotypes analyzed using SV-specific barcodes (not enough informative SNVs due to LOH). (*D*) Complex, intra-chromosomal rearrangement spanning approximately 0.5 Mb on 16q11.2 and 16q12.1 that involve two overlapping inversions, 63 kb (red) and 125 kb (blue), and a 163 kb tandem duplication (green). This rearrangement resides on the non-duplicated haplotype of this triploid region. *ORC6* is located entirely within the 63 kb inversion on 16q11.2 and is “deleted” the by the left breakpoint of the 125 kb inversion, which also inverts *MYLK3*. *C16orf8* on the same haplotype is also partially “deleted” by the 125 kb inversion (blue); *NETO2* is duplicated by the 163 kb tandem duplication (green). Inset: *MYLK3* and *ORC6* show allele-specific expression (Supplemental Table S12). *MYLK3* is only expressed from this rearranged allele (Haplotype 2); *ORC6* is expressed from the non-rearranged “diploid” allele (Haplotype 1).

In addition, we also leveraged the long-range information derived from the linked-reads to identify, assemble, and reconstruct SV-spanning breakpoints (including those of large-scale complex rearrangements) in the K562 genome using the recently established method Genome-wide Reconstruction of Complex Structural Variants (GROC-SVs) (Spies et al. 2017). In this method, long DNA fragments that span breakpoints are statistically inferred and refined by quantifying barcode similarity between pairs of genomic regions, similar to Long Ranger (Marks et al. 2018). Sequence reconstruction is then performed by assembling the relevant linked-reads around the identified breakpoints from which complex SVs are then automatically reconstructed. The breakpoints that have supporting evidence from the K562 3 kb-mate-pair dataset (see Supplemental Methods) were determined as high-confidence events (Dataset S5). GROC-SVs identified a total of 161 high-confidence breakpoints including 12 inter-chromosomal events (Fig. 1, Dataset S5, Fig. 4B); each event is accompanied with visualization (Supplemental Data); 138 of the breakpoints were successfully sequence-assembled with nucleotide-level resolution of breakpoints as well the exact sequence in the cases where nucleotides have been added or deleted (Dataset S5). A notable example of assembly by GROC-SVs is a complex intra-chromosomal rearrangement on chromosome 13 (Figure 4B).

Using the methods (“gemtools”) as described in (Greer et al. 2017), we identified phased structural rearrangements (multiple deletions and tandem duplications) within the tumor suppressor gene *FHIT* on 3p14.2 (Waters et al. 2014) (Fig. 4C). Since K562 exhibits LOH on chromosome 3, the SVs within *FHIT* were phased using linked-read barcodes instead of heterozygous SNVs. The hemizygous deletion between (59.74 Mb - 60.08 Mb) of 3p14.2 results in the loss of *FHIT* exons 6, 7, and 8. For the two phased tandem duplications on the same allele, one is intronic, and the other duplicates exon 5 (Fig. 4C). The two deletions downstream to the phased duplications are on two different alleles of *FHIT*. Another allele-specific, complex, intra-chromosomal rearrangement in K562 spans approximately 0.5 Mb on 16q11.2 and 16q12.1 (Fig. 4D), involving two overlapping inversions (62 kb and 125 kb) and a tandem duplication (163 kb). These events affect *ORC6*, *MYLK3*, *RHBDF1* also known as *C16orf8*, and *NETO2*, which has recently been identified as a cancer marker gene (Oparina et al. 2012; Hu et al. 2015). This rearrangement resides on the non-duplicated haplotype of this triploid region. *ORC6* is located entirely within the more centromeric inversion of this locus on 16q11.2 and is “deleted” the by left breakpoint of the more telemetric inversion, which also “deletes” *C16orf8* and inverts *MYLK3*, possibly disrupting its promoter region or proximal enhancers or disconnecting *MYLK3* from their regulation (Fig. 4D).

### Small-Scale Complex SVs from Deep-Coverage WGS

Small-scale complex SVs (Fig. 5A-E) as well as non-complex SVs were identified using a novel algorithm called Automated Reconstruction of Complex Structural Variants (ARC-SV) (Arthur et al. 2017) from deep-coverage WGS data (Dataset S6, Supplemental Data). These small-scale complex SVs are defined as genomic rearrangements with multiple breakpoints that cannot be explained by one well-defined (non-complex) SV type such as deletions, insertions, tandem duplications, or inversions. After filtering out SVs <50 bp or with breakpoints that reside in simple repeats, low complexity regions, satellite repeats, or segmental duplications, we identified 122 complex SVs (accompanied with schematic visualizations), 2,235 deletions, 320 tandem duplications, and 6 inversions (Dataset S6). Examples of complex SVs include dispersed duplications where duplicated sequences are inserted elsewhere in the genome in a non-tandem fashion (Fig. 5A). These dispersed duplications sometimes involve inversions of the inserted sequence and deletions at the insertion site (Fig. 5B, C). Other examples include inversions flanked on one or both sides by deletions (Fig. 5D), duplications that involve multiple non-exact copies, as well as deletion, inversion, and multiple duplications residing at the same locus (Fig. 5E). No other published algorithm to date has the capability to automatically identify and reconstruct these complex SVs. Eight out of ten breakpoints from five complex SVs were successfully validated by PCR and Sanger sequencing (Supplemental Table S8).

**Figure 5.**
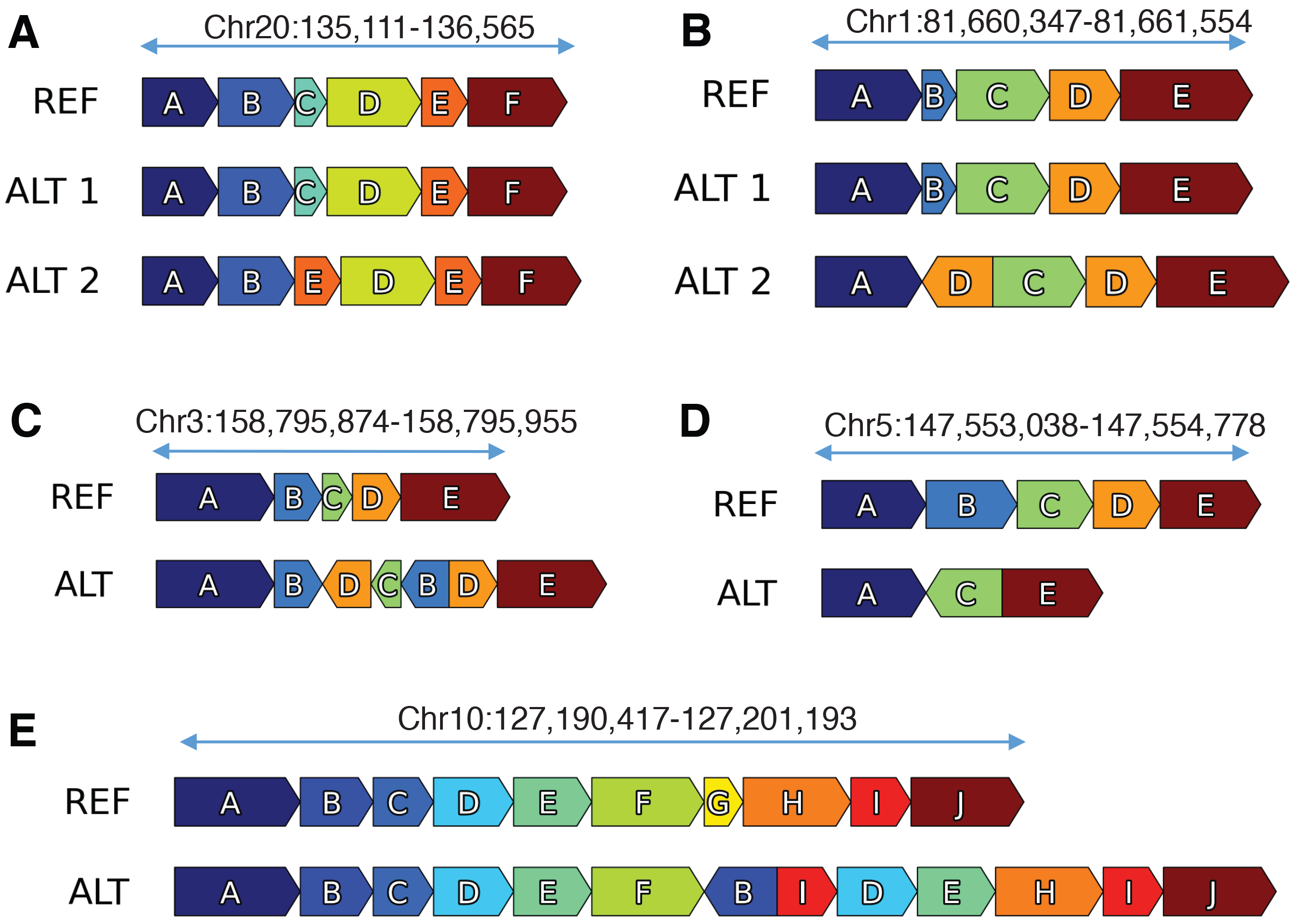
Small-scale complex SVs in K562 resolved using ARC-SV. Examples of small-scale complex SVs resolved using ARC-SV (Arthur et al. 2017) from the K562 WGS dataset. (*A*) Deletion of Block C and duplication of Block E between Blocks B and D on chromosome 20 (135,111-136,565). This variant has been validated by PCR. (*B*) Deletion of Block B and inverted duplication of Block D between Blocks A and C on chromosome 1 (81,660,347-81,661,554). (*C*) Duplication and inversion of Blocks B, C, and D between Blocks B and D on chromosome 3 (158,795,874-158,795,955) overlapping *IQCJ-SCHIP1*. (*D*) Inversion of Block C flanked by deletions of Blocks C and D on chromosome 5 (147,553,038-147,554,778) inside *SPINK14* (coding for a serine peptidase inhibitor). (*E*) Deletion of Block G, duplications of blocks I, D, and E, and inverted duplication of Block B between Blocks F and H on chromosome 10 (127,190,417-127,201,193).

### SVs from Mate-Pair Sequencing Analysis

To increase the sensitivity of detecting medium-sized SVs (1 kb-100 kb) in K562, we constructed a 3 kb-mate-pair library and sequenced (2 × 151 bp) to 6.9× non-duplicate coverage. The sequence coverage (C_R_) of each 3 kb insert is 302bp or 10%, which translates to a physical coverage (C_F_) of 68.5×. From the mate-pair library, SVs (deletions, inversions, and tandem duplications) were identified by clustering discordant read-pairs and split-reads using LUMPY (Layer et al. 2014). Only SVs that have both discordant read-pair and split-read support were retained. Overall, we identified 270 deletions, 35 inversions, and 124 tandem duplications using this approach (Dataset S7). Approximately 83% of these SVs are between 1 kb-10 kb, and 88% are between 1 kb-100 kb (Dataset S7). Twelve deletions and five tandem duplications were randomly selected for PCR and Sanger sequencing validation (Supplemental Table S8). The validation rates were 83% and 80% respectively.

### Non-Complex SVs from Deep-Coverage WGS

Non-complex SVs (deletions, inversions, insertions, and tandem duplications) in K562 were called from deep-coverage WGS data using a combination of established methods, namely Pindel (Ye et al. 2009), BreakDancer (Chen et al. 2009), and BreakSeq (Lam et al. 2010). These SVs were combined with those of the same SV type that were identified using ARC-SV, LUMPY and Long Ranger, where SVs (*n*=2,665) with support from multiple methods by ≥50% reciprocal overlap were merged. Through this combination of methods, a total of 9,082 non-complex SVs were identified in the K562 genome, including 5,490 deletions, 531 duplications, 436 inversions, and 2,602 insertions (Supplemental Data). (We note that only BreakDancer (Chen et al. 2009) was designed to call insertions.) Consistent with previous analyses (e.g. as in (Lam et al. 2012)), deletions show the highest number of concordant calls across the various methods compared to duplications and inversions ( Supplemental Fig S5, Supplemental Data). Eighteen deletions (>1 kb) and and 18 tandem duplications, both with split-read support, were randomly chosen for experimental validation using PCR and Sanger sequencing. The validation rates were 89% and 72% respectively ( Supplemental Table S8).

### LINE1 and Alu Insertions

We identified non-reference LINE1 and Alu retrotransposon insertions (REIs) in the K562 genome from our deep-coverage short-insert WGS data using a modified RetroSeq (Keane et al. 2013) approach (Supplemental Methods). Non-reference REIs were identified from paired-end reads that have one of the paired reads mapping to the human reference genome and the other read mapping to either the Alu or LINE1 consensus sequence in a full or split-read fashion (see Methods). We identified 1,147 non-reference Alu insertions and 85 non-reference LINE1 insertions in K562 (Supplemental Table S9, Fig. 1). Nine Alu and ten LINE1 insertions with split-read support were randomly chosen for validation using PCR and Sanger sequencing. The validation rates were 88% and 100% respectively (Supplemental Table S10). PCR primers were designed such that one anneals within the retrotransposon sequence and the other anneals in the unique sequences surrounding the predicted insertion site.

### Allele-Specific Gene Expression

Integrating CN information (i.e. allele frequencies) of the heterozygous SNVs (Dataset S1), we re-analyzed two replicates of ENCODE polyA-mRNA RNA-seq data to identify allele-specific gene expression in K562. We identified 5,053 and 5,149 genes that show allele-specific expression (*p* < 0.05) in replicates one and two, respectively (Fig. 1, Supplemental Table S11). We also identified 2,342 and 2,176 genes that would have been falsely identified to have allele-specific expression and 1,641 and 1,710 genes that would not have been identified to have allele-specific expression in replicates one and two, respectively, if the allele frequencies of heterozygous SNVs in aneuploid regions were not taken into consideration (Supplemental Table S12).

### Allele-Specific DNA methylation

By integrating CN and phase information of heterozygous SNVs of K562, we identified 110 CpG islands (CGIs) that exhibit allele-specific DNA methylation (Figure 1, Supplementary Table S13). We obtained K562 whole-genome bisulfite sequencing (WGBS) reads (2 × 100 bp, library ENCLB742NWU) from the ENCODE portal (Sloan et al. 2016) and aligned the reads to hg19 using Bismark (Krueger and Andrews 2011), where 76.9% of reads were uniquely mapped and 26.2% of cytosines were methylated in a CpG context. We then used reads that overlap both phased heterozygous SNVs (Dataset S2) and CpGs to phase the methylated and unmethylated CpGs to their respective haplotypes. We then grouped the phased individual CpGs into CGIs. Fisher’s exact test (taking the CN of a given CGI locus into account) was used to evaluate allele-specific methylation (*p*<0.05), and significant results were selected using a target false discovery rate of 10% (Supplemental Methods). Of these 110 CGIsZZ, 35 reside within promoter regions (here defined as 1 kb upstream of a gene); 83 are intragenic, and 28 lie within 1 kb downstream of 113 different genes. The following 6 genes are within 1 kb of a differentially methylated CGI and overlap with the Sanger Cancer Gene Census: *ABL1*, *AXIN2*, *CCND1*, *HOXD11*, *KDR*, and *PRDM16*.

### Allele-Specific CRISPR Targets

We identified a total of 28,511 targets in the K562 genome suitable for allele-specific CRISPR targeting (Fig. 1, Supplemental Table S14). Sequences (including reverse complement) of phased variants that differ by more than one base pair between the alleles were extracted to find all possible CRISPR targets by searching for the pattern [G, C, or A]N_20_GG (Supplemental Methods). Using a selection method previously described and validated (Sunagawa et al. 2016), only conserved high-quality targets were retained. We also took gRNA function and structure into consideration and performed furthering filtering of CRISPR targets. Targets with multiple exact matches, extreme GC content, and those containing TTTT (which might break the secondary structure of gRNA), were removed. We also used the Vienna RNA-fold package (Lorenz et al. 2011) to compute gRNA secondary structure and eliminated all targets for which the stem loop structure (for Cas9 recognition) could not form (Nishimasu et al. 2014). Finally, we calculated the off-target risk score by using the tool as described in (Ran et al. 2013). To ensure that all targets are as reliable and as specific as possible, we chose a very strict threshold and rejected candidates with a score below 75. Of the 28,511 allele-specific CRISPR target sites, 15,488 are within an annotated protein-coding or non-coding RNA transcript, 705 within an exon, and 13 targets are within an experimentally validated enhancer (Visel et al. 2007) (Supplementary Table S14).

### Genomic Structural Context Provides Insight into Regulatory Complexity

We show examples of how deeper insights into gene regulation and regulatory complexity can be obtained by integrating genomic structural contexts with functional genomics and epigenomics data (Fig. 6A-D). One example is the allele-specific RNA expression and allele-specific DNA methylation in K562 at the *HOXB7* locus on chromosome 17 (Fig. 6A). By incorporating the genomic context of *HOXB7* in K562, we see that *HOXB7* exhibit highly preferential RNA expression from the two copies of Haplotype 1 (*p* = 0.007) in which the CGI near its promoter is completely unmethylated (*p* = 3.18 × 10^−18^) (Fig. 6 A, C). The second example is allele-specific RNA expression and allele-specific DNA methylation of the *HLX* gene in K562 (Fig. 6B). The *HLX* locus on chromosome 1 is tetraploid, and we see that *HLX* is only expressed from Haplotype 1 which has three copies and not expressed in Haplotype 2 (*p* = 0.043) (Fig. 6B, D). The CGI of the *HLX* locus is unmethylated in Haplotype 2 but highly methylated on Haplotype 1 (*p* = 5.14 × 10^−15^) (Fig. 6B, C). There is also an allele-specific CRISPR targeting site for both haplotypes within *HLX* (Fig. 6B). In addition, we performed Pearson correlation analysis between our deep-coverage K562 WGS data and K562 POLR2A ChIP-seq data (previously released on the ENCODE data portal) to determine whether changes in K562 genome CN or ploidy affected binding of the polymerase molecule to genomic DNA in a large-scale fashion (Supplemental Fig. S6). The two sets of data are very well correlated (*r*=0.51, *p*<2.2 × 10^−16^) suggesting that RNA polymerase activity is generally influenced by ploidy in the K562 genome. In addition, we also correlated the K562 POLR2A ChIP-seq data with the FPKM values from four independent K562 polyA RNA-seq experiments (also previously released on the ENCODE portal) and find that these datasets are also very well correlated consistently (*r*=0.46, *p*<2.2 × 10^−16^; *r*=0.58, *p*<2.2 × 10^−16^; *r*=0.47, *p*<2.2 × 10^−16^; *r*=0.46, *p*<2.2 × 10^−16^) (Supplemental Fig. S7A-D).

**Figure 6.**
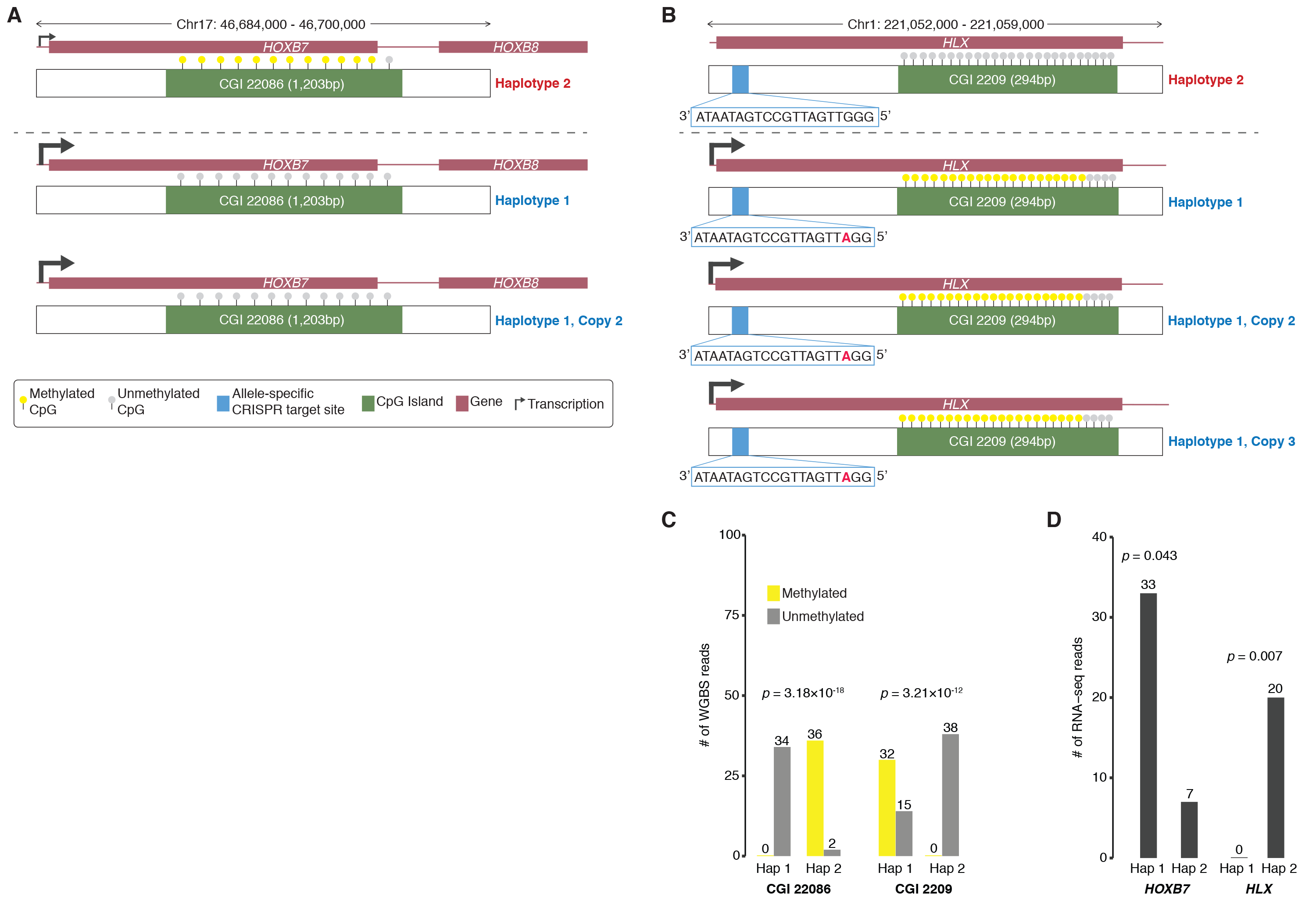
Genomic structural contexts provide insights into regulatory complexity. (*A*) Chr17:46,687,000-46,700,000 locus (triploid in K562) containing *HOXB7* and *HOXB8* and CpG Island (CGI) 22086 (1,203 bp) where phased Haplotype 1 has two copies and Haplotype 2 has one copy. Allele-specific expression of *HOXB7* from Haplotype 1. CpGs in CGI 22086 are unmethylated in Haplotype 1 and methylated in Haplotype 2. (*B*) Chr1:221,052,000-221,059,000 locus (tetraploid in K562) containing *HLX* and CGI 2209 (294 bp) where phased Haplotype 1 has three copies and Haplotype 2 has one copy. Allele-specific expression of *HLX* from Haplotype 1. CpGs in cGi 2209 are unmethylated in Haplotype 2 and highly methylated in Haplotype 1. Allele-specific CRISPR targeting site 797 bp inside the 5’ end of the *HLX* for both Haplotypes. (*C*) Number of methylated and unmethylated phased WGBS reads for Haplotypes 1 and 2 in CGI 22086 and CGI 2209 where both CGIs exhibit allele-specific DNA methylation. (*D*) Number of RNA-seq reads for Haplotypes 1 and 2 of *HLX* and *HOXB7* where both genes exhibit allele-specific RNA expression.

Furthermore, we also find allele-specific RNA expression for the rearranged copy of *MYLK3* (*p*<1.93 × 10^−17^) and the normal, non-rearranged copies (CN=2) of *ORC6* (*p*<1.58 × 10^−8^) where expression from re-arranged allele (CN=1) of *ORC6* is “depleted” in K562 (Table S11, Fig. 4C, D). These observations made by integrating our K562 linked-read data and ENCODE RNA expression data provide novel insights into gene regulatory mechanisms in terms of ectopic expression and dosage compensation, which also raises important questions regarding the history of the K562 cell line in terms of mutations, selective pressures, and adaption (see Discussion).

## DISCUSSION

K562 is one of the most widely used laboratory “work-horse” cell lines in the world. Among the three tier-one cell lines of ENCODE, K562 has by far the most functional genomics and epigenomics data generated. Furthermore, K562 is also one of the most commonly used cell lines for large-scale CRISPR/Cas9 gene-targeting screens (Wang et al. 2015; Arroyo et al. 2016; Morgens et al. 2016; Han et al. 2017; Adamson et al. 2016; Liu et al. 2017). Yet, despite its wide usage and impact on biomedical research, its genomic sequence and structural features have never been comprehensively characterized, beyond its karyotype (Selden et al. 1983; Gribble et al. 2000; Wu et al. 1995; Naumann et al. 2001) and SNPs called from 30×-coverage WGS but without taking aneuploidy or CN into consideration (Cavalli et al. 2016). Analysis, integration, and interpretation of the extensive collection of functional genomics and epigenomics datasets for K562 had so far relied solely on the human reference genome. Here, we present the first detailed and comprehensive characterization of the K562 genome so that future studies no longer have to rely solely on the human reference genome. By performing deep-coverage short-insert WGS, 3 kb-insert mate-pair sequencing, deep-coverage linked-reads sequencing, array CGH, karyotyping, and integrating a compendium of novel and established analysis methods (Supplemental Fig. S1A), we produced a comprehensive spectrum of genomic structural features (Fig. 1) for K562 that includes SNVs (Dataset S1), Indels (Dataset S1), ploidy by chromosome segments at 10-kb resolution (Supplemental Table S2), phased haplotypes (Dataset S2, Supplemental Data)—often of entire chromosome arms (Table 2, Supplemental Data)—phased CRISPR targets (Supplemental Table S14), nonreference retrotransposon insertions (Table S9), and SVs (Supplemental Data) including deletions, duplications, and inversions, and complex SVs (Dataset S6, Dataset S7). Many SVs were also phased, assembled, and experimentally verified (Dataset S2-S5, Supplemental Table S8, Supplemental Table S10). Of the 3,784,863 variants that were haplotype-phased in the K562 genome (Dataset S2-S5), 3,088,185 (81.6%) are SNVs; 692,998 (18.31%) are Indels; 3,451 are deletion SVs (51 bp to 30 kb) (0.1%), and 229 are large SVs.

Pervasive aneuploidy is a characteristic of many cancers. Previous studies have confirmed the near triploid karyotype of K562 (Selden et al. 1983; Gribble et al. 2000; Wu et al. 1995; Naumann et al. 2001). In our analysis, however, we also found considerable portions of the K562 genome to be much more varied than what had previously been reported. This is because by leveraging deep-coverage WGS, the CN across different chromosome segments, as determined by our read-depth analysis, is of much higher resolution than karyotyping. Furthermore, the identified chromosome segments with aneuploidy (CN>2) and orthogonally supported by karyotyping and array CGH were further validated, also orthogonally, from a statistical approach in which significant differences in unique linked-read barcode counts between the major and minor haplotypes were determined using a one-sided *t*-test (p < 0.001) (Bell et al. 2017). In addition, it has to be taken into consideration that for a widely used cell line with decades of history such as K562, additional genome variation is expected. In light of this, it is reassuring that the overall karyotype has not changed much over the years. However, researchers should still keep this aspect in mind when working with a version of K562 that has been separated from the main ENCODE K562 line used here. We expect that the vast majority of genomic variants that we describe here to be universal for K562, but for individual variants, it is possible that different lines of K562 may have slightly diverged from each other (Supplemental Table S1, Supplemental Table S3). When using a different K562 line and following up on findings for individual loci of interest, a first step should always be to experimentally validate their presence. When incorporating these genomic variants for global analyses, such as interrogating network interactions, the vast majority of them will exist, thus such global analyses are expected to yield substantial insights. Even though the pervasive aneuploidy in K562 renders the design and interpretation of K562 studies more challenging, the information we provide here enables researchers to continue the use of this cell line to investigate the effects of genetic variation on the multiple levels of functional genomics activity and regulation for which ENCODE data already exists or continues to be produced. Thus, analysis of K562 data should not only be more complex and challenging, but also potentially much more insightful and rewarding when taking its complex genome structure into account.

Sensitive and accurate identification of SNVs and Indels requires relatively deep WGS coverage (>33× and >60× respectively) (Bentley et al. 2008; Fang et al. 2014). From our >70× non-duplicate coverage WGS data, we identified large numbers of SNVs and Indels that we could subsequently correct for their allele frequencies according to ploidy. In addition to being essential for correct haplotype identification, these ploidy-corrected variants are also needed for functional genomics or epigenomics analyses such as the determination of allele-specific gene expression or of allele-specific transcription factor binding in K562 (Cavalli et al. 2016). From RNA-seq or ChIP-seq data analysis, a statistically significant increase in transcription or transcription-factor-binding signal in one allele compared to the other at a heterozygous locus, may be identified as a case of allele-specific expression or allele-specific transcription-factor binding which usually suggests allele-specific gene regulation at this locus. However, if aneuploidy can be taken into consideration and the signals normalized by ploidy, the case identified might be a result of increased CN rather than the preferential activation of one allele over the other on the epigenomic level. Indeed, in our re-analysis of two replicates of ENCODE K562 RNA-seq data, we identified 2,359 and 2,643 genes that would have been falsely identified to have allele-specific expression in addition to 1,808 and 2,063 genes that would not have been identified to have allele-specific expression in replicates one and two, respectively, if ploidy was not taken into consideration (Supplemental Table S12).

The haplotype phase of genomic sequence variants is an essential aspect of human genetics, but current standard WGS approaches entirely fail to resolve this aspect. We performed linked-read sequencing of K562 genomic DNA using the Chromium System from 10× Genomics (Zheng et al. 2016; Marks et al. 2018). After size-selecting for genomic DNA fragments >30kb, 300 genomic equivalents of HMW DNA were partitioned into more than one million oil droplets, uniquely barcoded within each droplet, and subjected to random priming and amplification. Implemented by Long Ranger (Marks et al. 2018), sequencing reads that originate from the same HMW DNA molecule can be identified by their respective droplet barcodes and linked together to produce virtual long reads. Then, by looking for virtual long reads that overlap a previously called set of heterozygous haplotypes (Dataset S1), the phase information of the heterozygous haplotypes was determined and the virtual long reads were constructed into phased haplotype blocks with N50 > 2.72 Mb (Dataset S2, Fig. 2D). Chromosomes 3, 9, 13, 14, X, and large portions of chromosomes 2, 10, 12, 20, 22 were difficult to phase, resulting in comparatively shorter phased blocks (Dataset S2, Fig. 1, Supplemental Fig. S4). This is not surprising since these chromosomes and chromosomal regions exhibit a very high degree of LOH (Fig. 1 and Supplemental Table S4). Heterozygous loci in aneuploidy regions with more than two haplotypes were excluded from phasing linked-read analysis due to software and algorithmic limitations (Zheng et al. 2016). However, the phase information of these loci could be resolved from our linked-read data in principle, should new algorithms become available.

We extended on the already-impressive phasing capabilities of Long Ranger and constructed mega-haplotypes in K562—often spanning entire chromosome arms (Fig. 3, Table 2, Supplementary Data)—by leveraging the haplotype imbalance in aneuploid chromosomes using a recently developed method for which its effectiveness in cancer has already been demonstrated (Bell et al. 2017). Since gene dosage is a fundamental component of genome biology and for which aneuploidy contribute large effects in terms of amplification and reduction, the ability to haplotype across long stretches of aneuploidy is essential for understanding of the genetic regulations of cancer and an important component for developing genetically targeted cancer treatment.

It has been shown previously that integrating orthogonal methods and signals improves SV-calling sensitivity and accuracy (Mohiyuddin et al. 2015; Layer et al. 2014). Here, we combined deep-coverage short-insert WGS, mate-pair sequencing, linked-read sequencing, and several SV-calling methods to identify many non-complex SVs. To obtain the union set of noncomplex SV calls from the various methods, the SVs identified by multiple methods were merged and indicated accordingly (Supplemental Data). For deletions (Supplemental Fig. S5A), we see strong overlap for the various methods, but this overlap is less pronounced for duplications (Supplemental Fig. S5B) and inversions (Supplemental Fig. S5C). This is consistent with previous analysis (Lam et al. 2012) as inversions and duplications are more difficult in principle to accurately resolve (Lin et al. 2015; Sudmant et al. 2015). We also expect the detection of many SVs to be method-specific, since each method is designed to utilize different types of signals and also optimized to identify different classes of SVs (Pabinger et al. 2014; Lin et al. 2015). Again, if particular SVs are of interest for follow-up studies, they should first be experimentally validated.

The complex rearrangements identified by using ARC-SV from short-insert WGS (Fig. 5A-E, Dataset S6, Supplemental Data) and by using GROC-SVs from linked-reads (Fig. 4B, Dataset S5, Supplemental Data) are classes of SVs that could not be easily identified and automatically reconstructed using previously existing methods. The small-scale complex SVs that were identified by using ARC-SV and experimentally validated (Fig. 5, Supplemental Table S8, Supplemental Data) describe a subtle class of complex rearrangements in cancer genomes that have never been previously demonstrated by others, but have long speculated to exist (Perry et al. 2008; Quinlan and Hall 2012; Collins et al. 2017). Detecting and automatically reconstructing these small-scale complex SVs, especially in a “hay” of canonical SVs and in highly rearranged cancer genomes, has remained an unsolved problem for many years. In other words, our results reveal a class of previously overlooked complex SVs in cancer that can now be identified from standard short-insert WGS data and elucidated further. They have clear implications for the conventional models of cancer evolution which often assume gradual, step-by-step mutations; however, these complex SVs support a form of punctuated genome evolution (Davis et al. 2017). A major unsolved question still is how complex SVs arise mechanistically for which there are general models: template switching during replication (Lee et al. 2007; Hastings et al. 2009) and chromothripsis (Stephens et al. 2011). These small-scale complex SVs in K562 implicate that another, yet-undescribed, mechanism might also be contributing, at least in cancer. Furthermore, the functional consequences of these small-scale complex SVs are also unknown. These important questions remain unsolved mainly due to the lack of data and examples. It is possible that this mutational complexity contributes to genome innovation, at least in cancer, or is just a curious sideshow (Quinlan and Hall 2012). Only the accumulation of such examples and data will allow researchers in fields such as cancer evolution to begin to address these important questions.

Before the existence of linked-read sequencing, haplotype phasing and resolving large SVs (>30kb) relied heavily on fosmid libraries (Snyder et al. 2015; Williams et al. 2012; Kitzman et al. 2011; Cao et al. 2015; Adey et al. 2013) which were laborious, costly, time consuming, and much less efficient. Using linked-read sequencing and gemtools (Greer et al. 2017), we phased and resolved complex SVs that are especially compelling on 3p14.2 (within the tumor suppressor gene *FHIT*) and on 16q11.2 and 16q12.1 of the K562 genome. *FHIT* is frequently seen to harbor deletions in many types of human cancers, most commonly of epithelial origin, such as lung, stomach, cervix, head and neck, breast, and kidney (Lubinski et al. 1994; Ohta et al. 1996; Huebner et al. 1998; Ingvarsson 2001). Reduction or absence in its protein expression occurs in nearly 50% of all cancers (Huebner et al. 1998; Waters et al. 2014). While LOH and allele-specific deletions within *FHIT* have been previously reported (Wistuba et al. 1997; Li et al. 2016), to our knowledge, this is the first discovery of phased and allele-specific tandem duplications within *FHIT* and the first report of *FHIT* mutations for CML. Curiously, all previous reports of deletions within *FHIT* for various cancer types (not including CML) were all centered on and include exon 5 (Durkin et al. 2008), whereas exon 5 is duplicated in K562. Deletions of all three *FHIT* exons 6, 7, and 8 (Fig. 4C) are less frequent but have been reported for lung cancer and esophageal adenocarcinoma (Sozzi et al. 1996; Dagmar et al. 1997).

We identified highly allele-specific RNA expression for *ORC6* (*p*<1.58 × 10^−8^) and *MYLK3* (*p*<1.93 × 10^−17^) in K562 (Supplemental Table S11), which is likely contributed by the allele-specific complex intra-chromosomal rearrangement residing on the non-duplicated haplotype of 16q11.2 (triploid) (Fig. 4D). *ORC6* codes for origin recognition protein complex subunit 6 and is essential for coordinating DNA replication, chromosome segregation, and cytokinesis (Prasanth et al. 2002). One allele of *ORC6* is “deleted” by one of the inversion breakpoints of this rearrangement and maintains allele-specific expression on the other allele (Fig. 4D). The origin recognition complex, in which Orc6 is a subunit, serves as a “landing pad” for bringing together components of the pre-replicative complex required for DNA replication. Reduction of Orc6 dosage by small interfering RNA results in decreased DNA replication, aberrant mitosis, and the formation of multiple nuclei and multipolar spindles in cells; a long period of this reduction increases cell death (Prasanth et al. 2002). Such effects are not observed in K562 cells even though RNA expression from one allele of *ORC6* is “depleted” by this rearrangement (*p*<1.58 × 10^−8^, Supplemental Table S11). This is likely because the other “normal” *ORC6* allele was duplicated, rendering this locus in K562 triploid and thus maintain a “diploid” gene dosage. Perhaps more importantly, this insight also raises important questions regarding the history of mutation as well as selective pressures that occurred within K562 cells. It is possible that one copy of chromosome 16 was first duplicated, freeing this locus from selective pressures, allowing it to acquire new mutations, since “diploid” copies are still maintained in the genome. It is also conceivable that duplication of this locus is disadvantageous for cell proliferation, and K562 cells that acquired this rearrangement after the duplication of chromosome 16 also acquired a selective advantage since they now reverted this locus back to “diploid” copies. However, it is also possible, though perhaps less likely, that this rearrangement occurred before chromosome 16 duplication, putting a negative selection pressure on K562 cells, and that this pressure is released by duplication of the other copy. Interestingly, this rearrangement also inverts an intact copy of *MYLK3* (Fig. 4D), which was identified to encode a novel cardiac-specific myosin light chain kinase (Seguchi et al. 2007). Since its expression is expected to be normally repressed except in heart cells, the allele-specific RNA expression of *MYLK3* (*p*<1.93E-17, Supplemental Table S11) from this inverted allele suggests that this inversion activated the ectopic expression of this gene in K562, possibly by disrupting or disconnecting it from its promoter or proximal enhancer elements that impose repressive regulatory mechanisms. Finally, this complex rearrangement on chromosome 16 of K562 also duplicates *NETO2* in a tandem fashion, also on the same allele (Fig. 4D). *NETO2* codes for a single-pass membrane protein neuropilin and tolloid-like 2 (Stöhr et al. 2002). Its expression is frequently up-regulated in many types of human cancers including lung, cervical, colon, and renal carcinomas and has been suggested as a potential genetic marker for cancer (Oparina et al. 2012). Its up-regulation also correlates with the progression and poor prognosis of colorectal carcinoma (Hu et al. 2015). It is conceivable that the increase in *NETO2* gene dosage due to duplication in K562 cells contributes to their efficient proliferation in culture and that, at least in some cancers, the frequent up-regulations observed for *NETO2* are also contributed by this similar mechanism of allele-specific tandem duplication.

The hallmark of CML is the Philadelphia rearrangement t(9; 22)(q34; q11) which results in the fusion of *ABL1* and *BCR* (Heisterkamp et al. 1985; de Klein et al. 1982; Groffen et al. 1984). This gene fusion is known to be extensively amplified in the K562 genome by tandem duplication (Wu et al. 1995). FISH analysis showed that fluorescent signals from the *BCR/ABL1* gene fusion almost always concentrate on a single marker chromosome (Tkachuk et al. 1990; Wu et al. 1995; Gribble et al. 2000). This is also consistent with our data as the linked-reads that support the *BCR/ABL1* gene fusion do not share overlapping barcodes with linked-reads that align elsewhere in the genome, and the *BCR* and *ABL1* gene regions where the fusion occurs show a >2.8× increase in sequencing coverage relative to average sequencing coverage across the genome.

Data generated from this comprehensive whole-genome analysis of K562 is available through the ENCODE portal (Sloan et al. 2016) (Supplemental Figure S1B, C). We envision that this analysis will serve as a valuable resource for further understanding the vast troves of ENCODE data available for K562, such as determining whether a potential or known regulatory sequence element has been altered by SNVs or SNPs, Indels, retrotransposon insertions, a gain or loss of copies of that given element, or allele-specific regulation. As additional examples of how integrating genomic context can yield further understanding of existing ENCODE data, we showed, as examples, the complex gene regulatory scenarios uncovered at the *HOXB7* and *HLX* loci in K562. Hox genes are known to have important roles in hematopoiesis and oncogenesis (Argiropoulos and Humphries 2007; Shah and Sukumar 2010; Eklund 2011). The HOXB7 transcription factor mediates lymphoid development, hematopoietic differentiation and leukemogenesis (Giampaolo et al. 1995; Carè et al. 1999). *HOXB7* overexpression has been reported in leukemia (Raval et al. 2007) as well as in many other cancers (Caré et al. 1996; Wu et al. 2006; Yamashita et al. 2006; Shiraishi et al. 2007; Chen et al. 2008; Storti et al. 2011). It is directly upstream of *HOXB8*, which is the first Hox gene found to be an oncogene in leukemia (Blatt et al. 1988). *HLX* has also been suggested to play oncogenic roles in leukemia (Deguchi et al. 1992; Deguchi and Kehrl 1993; Jawad et al. 2006; Fröhling 2012). By integrating the genomic context of *HOXB7* and *HLX* in K562 with RNA-seq and WGBS data, we see that the RNA of both genes are expressed from haplotypes that exhibit aneuploidy and in an allele-specific manner (Fig. 6A, B, D). The allele-specific methylation of the CGIs near these two genes is associated with active transcription in the case of *HLX* and silencing of transcription in the case of *HOXB7* (Fig. 6A-C). Such insights into potential oncogene regulation cannot be obtained by analyzing functional genomics and epigenomics data alone without genome structural information i.e. correct genomic context. In addition, we also observed that the K562 POLR2A ChIP-seq signal in both replicates is very well correlated with polyA RNA-seq signal and with WGS coverage, suggesting an association between polymerase binding and active transcription and between polymerase binding and ploidy (Supplemental Figures S6, S7).

Our work here serves to guide future studies that utilize the K562 “workhorse” cell line, such as CRISPR screens where knowledge of the sequence variants can extend or modify the number of editing targets (Table S13) while knowledge of aberrant CN will allow for much more confident data interpretation. To give an example, in a recent study that uses CRISPRi to screen and elucidate the function of long non-coding RNAs in human cells, out of the seven cell types studied, the number of gRNA hits varied considerably among the various cell types, with 89.4% of hits unique to only one cell type and none in more than five cell types (Liu et al. 2017). Although a large portion of the phenomenon are very likely explained by cell-specific effects, it is still quite possible that many of the gRNA hit differences were the result of differences in genome sequence or ploidy. Our list of allele-specific CRISPR targets (Table S13) will allow for the discernment between these two potential reasons for differences in CRISPR effects and should be particularly valuable for future large-scale CRISPR screens that utilize K562. Lastly, this study also serves as a technical example for the advanced, integrated, and comprehensive analyses of other heavily utilized cell lines and genomes in biomedical research such as HepG2.

## METHODS

### Overview

We combined multiple experimental and analysis methods (Supplementary Fig. S1A), including karyotyping, array CGH, deep (72× non-duplicate coverage) short-insert whole-genome sequencing (WGS), 3 kb-mate-pair sequencing (Korbel et al. 2007) and 10x Genomics linked-reads sequencing (Zheng et al. 2016; Marks et al. 2018), to comprehensively characterize the genome of the primary ENCODE cell line K562 (Fig. 1). The WGS dataset was used to identify CN i.e. ploidy by chromosome segments, SNVs, Indels, non-reference LINE1 and Alu insertions (Lupski 2010; Sudmant et al. 2015), and SVs such as deletions, duplications, inversions, insertions, and small-scale complex SVs. These SVs were identified using an integrated approach that includes BreakDancer (Chen et al. 2009), Pindel (Ye et al. 2009), BreakSeq (Lam et al. 2010) and ARC-SV (Arthur et al. 2017). The allele frequencies of heterozygous SNVs and Indels were determined by taking ploidy into account. The linked-reads were used to phase SNVs and Indels as well as to identify, phase, reconstruct, and assemble primarily large (>30 kb) and complex SVs (Greer et al. 2017; Spies et al. 2017; Marks et al. 2018), though additional small-scale deletions were also identified and phased (Zheng et al. 2016). SVs and REIs were experimentally validated with PCR and Sanger sequencing. Phased SNV haplotype blocks in aneuploid regions were “stitched” to mega-haplotypes by leveraging haplotype imbalance (Bell et al. 2017). The 3 kb-mate-pair data was used to identify additional SVs and was also used to validate large and complex SVs identified from linked-reads. Functional genomics and epigenomics datasets from ENCODE were integrated with CN and phasing information to identify allele-specific RNA expression and allele-specific DNA methylation. Phased variants were also used to identify allele-specific CRISPR targets in the K562 genome. For full descriptions of experimental and computational procedures (including analysis code), see Supplemental Methods.

## DATA ACCESS

Raw and processed data generated in this study are publicly available on the ENCODE portal (encodeproject.org) (Sloan et al. 2016) under experiment accessions: ENCSR711UNY, ENCSR025GPQ, and ENCSR053AXS. ENCODE accessions for individual data files including Datasets S1-S7 are listed in Supplemental Fig. 1B, C. Analysis code is provided in Supplemental Methods. Overview of all resources generated for K562 in this study is listed with detailed descriptions in Supplemental Fig. 1B-D.

## ACKNOWLEDGEMENTS

We thank Aditi Narayanan, Dr. Idan Gabdank, Nathaniel Watson, Dr. Carrie Davis, Kathrina Onate, and Dr. Cricket Sloan for assistance with data upload to the ENCODE portal. We thank Dr. Athena Cherry and the Stanford Cytogenetics Laboratory for karyotype analysis and Arineh Khechaduri for performing genomic DNA preparation. We thank Dr. Minyi Shi for providing K562 cells. A.E.U. was supported by NIH grant P50-HG007735 and the Stanford Medicine Faculty Innovation Program, and B.Z. was additionally supported by NIH training grant T32-HL110952. W.H.W. received support from NIH grants HG007834 and HG007735. J.G.A. received funding from NIH training grant T32-GM096982 and NSF Graduate Fellowship DGE-114747. A.A. was funded by NIH grant U24CA220242.

## AUTHOR CONTRIBUTIONS

B. Z. and A.E.U conceived and designed the study. B.Z., R.P., N.B.E, M.S.H, and R.R.H performed experiments. B.Z., S.S.H., S.U.G., J.M.B., N.S., XW.Z., XL.Z., S.B., J.G.A., G.S., D.P., and A.A. performed analysis. H.P.J, W.H.W., and A.E.U contributed resources and supervised the study. B.Z., S.S.H., and A.E.U. wrote the manuscript.

---

**Table 1. Summary of K562 SNVs and Indels**

**Table 2. Haplotypes constructed in aneuploid regions by leveraging haplotype imbalance**

